# Structural dynamics of the SARS-CoV-2 frameshift-stimulatory pseudoknot reveal topologically distinct conformers

**DOI:** 10.1101/2020.12.28.424630

**Authors:** Krishna Neupane, Meng Zhao, Aaron Lyons, Sneha Munshi, Sandaru M. Ileperuma, Dustin B. Ritchie, Noel Q. Hoffer, Abhishek Narayan, Michael T. Woodside

## Abstract

The RNA pseudoknot that stimulates −1 programmed ribosomal frameshifting in SARS coronavirus-2 (SARS-CoV-2) is a possible drug target. To understand how this 3-stemmed pseudoknot responds to the mechanical tension applied by ribosomes during translation, which is thought to play a key role during frame-shifting, we probed its structural dynamics under tension using optical tweezers. Unfolding curves revealed that the frameshift signal formed multiple different structures: at least two distinct pseudoknotted conformers with different unfolding forces and energy barriers, as well as alternative stem-loop structures. Refolding curves showed that stem 1 formed first in the pseudoknotted conformers, followed by stem 3 and then stem 2. By extending the handle holding the RNA to occlude the 5′ end of stem 1, the proportion of the different pseudoknot conformers could be altered systematically, consistent with structures observed in cryo-EM images and computational simulations that had distinct topologies: the 5′ end of the RNA threaded through the 3-helix junction to form a ring-knot, or unthreaded as in more standard H-type pseudoknots. These results resolve the folding mechanism of the frameshift signal in SARS-CoV-2 and highlight the dynamic conformational heterogeneity of this RNA, with important implications for structure-based drug-discovery efforts.

---

Like most coronaviruses, the Severe Acute Respiratory Syndrome coronavirus 2 (SARS-CoV-2) causing the COVID-19 pandemic makes use of −1 programmed ribosomal frameshifting (−1 PRF) to express proteins that are essential for viral replication^1^. In −1 PRF, a shift in the reading frame of the ribosome at a specific location in the RNA message is stimulated by a structure in the mRNA located 5–7 nt downstream of the ‘slippery’ sequence where the reading-frame shift occurs, thereby generating alternate gene products^2,3^. Previous work on viruses including HIV-1 and SARS-CoV showed that mutations modulating the level of −1 PRF can significantly attenuate viral propagation in cell culture^4–6^. As a result, the structures stimulating −1 PRF are potential targets for anti-viral drugs^7–9^, motivating efforts to find ligands active against −1 PRF in SARS-CoV-2 that could be used to treat COVID-19 ^10–14^.

The pseudoknot stimulating −1 PRF in SARS-CoV-2 has a 3-stem architecture^1,10,15,16^ (Fig. 1A) that is characteristic of coronaviruses, in contrast to the more common 2-stem architecture of most viral frameshift-stimulatory pseudoknots^17^. Cryo-EM imaging^10,15^ and computational modeling^18^ both suggest that the SARS-CoV-2 pseudoknot can take on several different conformers (Fig. 1B, C). Some of these conformers involve knot-like fold topologies that have not previously been observed in frameshift-stimulatory pseudoknots, specifically conformers with the 5′ end threaded through the junction between the 3 helices to generate what we term a ‘ring-knot’^10,15,18^. Such a 5′-end threaded ring-knot fold has only previously been observed in viral exoribonucle-ase-resistant RNAs^19–21^. Intriguingly, the co-existence of multiple conformers in the SARS-CoV-2 pseudoknot is consistent with evidence from studies of various stimulatory structures, both pseudoknots and hairpins^22–25^, as well as from studies of the effects of anti-frameshifting ligands^26^, which shows that the stimulation of −1 PRF is linked primarily to conformational heterogeneity in the stimulatory structure. In particular, −1 PRF is linked to the conformational heterogeneity under tension^27^ in the range of forces applied by the ribosome during translation^28,29^. However, the dynamic ensemble of conformers populated by the SARS-CoV-2 pseudoknot has not yet been explored experimentally, and the folding mechanism of this pseudoknot—especially its unusual ring-knotted conformer—remains unknown.

**Fig. 1:**
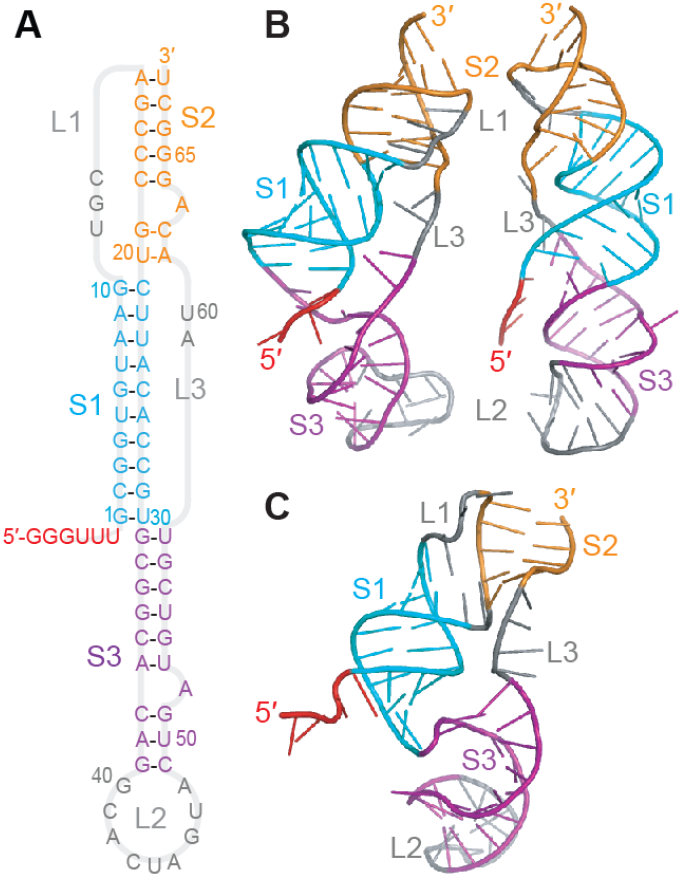
3-stemmed architecture of SARS-CoV-2 frameshift-stimulatory pseudoknot. (A) Secondary structure from Ref. 1, with stems and loops color-coded. (B) 3D structures of 5′- threaded (left) and unthreaded (right) conformers from Ref. 18. (C) Cryo-EM structure from Ref. 15.

Here we examine the conformational dynamics of the SARS-CoV-2 pseudoknot in the single-molecule regime. We study it under tension in optical tweezers in order to mimic the situation seen during −1 PRF, where the force applied by the ribosome is ramped up and down as the ribosome attempts to resolve the mRNA structure before shifting reading frame^30^. Such force spectroscopy measurements are also a powerful tool for characterizing folding mechanisms^31^ and the energy landscapes that govern folding dynamics^32^. We find that the SARS-CoV-2 frameshift signal indeed forms at least 2 distinct pseudoknotted conformers, one involving threading of the 5′-end to form a ring-knot and the other without any threading. Stem 1 usually folds first, followed by stem 3 and lastly stem 2, but sometimes alternate stem-loops form that displace the pseudoknotted structures. The existence of multiple conformers of this frameshift signal has important implications for structure-based efforts to find small-molecule therapeutics targeting −1 PRF.

## Results

To probe the conformations formed by the SARS-CoV-2 pseudoknot, their folding pathways, and the dynamics under tension, we annealed a single RNA molecule containing the sequence of the pseudoknot flanked by ‘handle’ regions to DNA handles that were attached to beads held in optical traps (Fig 2A). We then moved the traps apart to ramp up the force and unfold the RNA, and brought them back together to ramp down the force and refold the RNA. Force-extension curves (FECs) measured during unfolding in near-physiological ionic conditions (130 mM K^+^, 4 mM Mg^2+^) showed one or more characteristic transitions in which the extension abruptly increased and force simultaneously decreased when part or all of the structure unfolded cooperatively (Fig. 2B). Unfolding events were observed over a range of forces from ~5–50 pN; similar transitions were seen in refolding FECs, but at forces below ~15 pN (Fig. 2C).

**Fig. 2:**
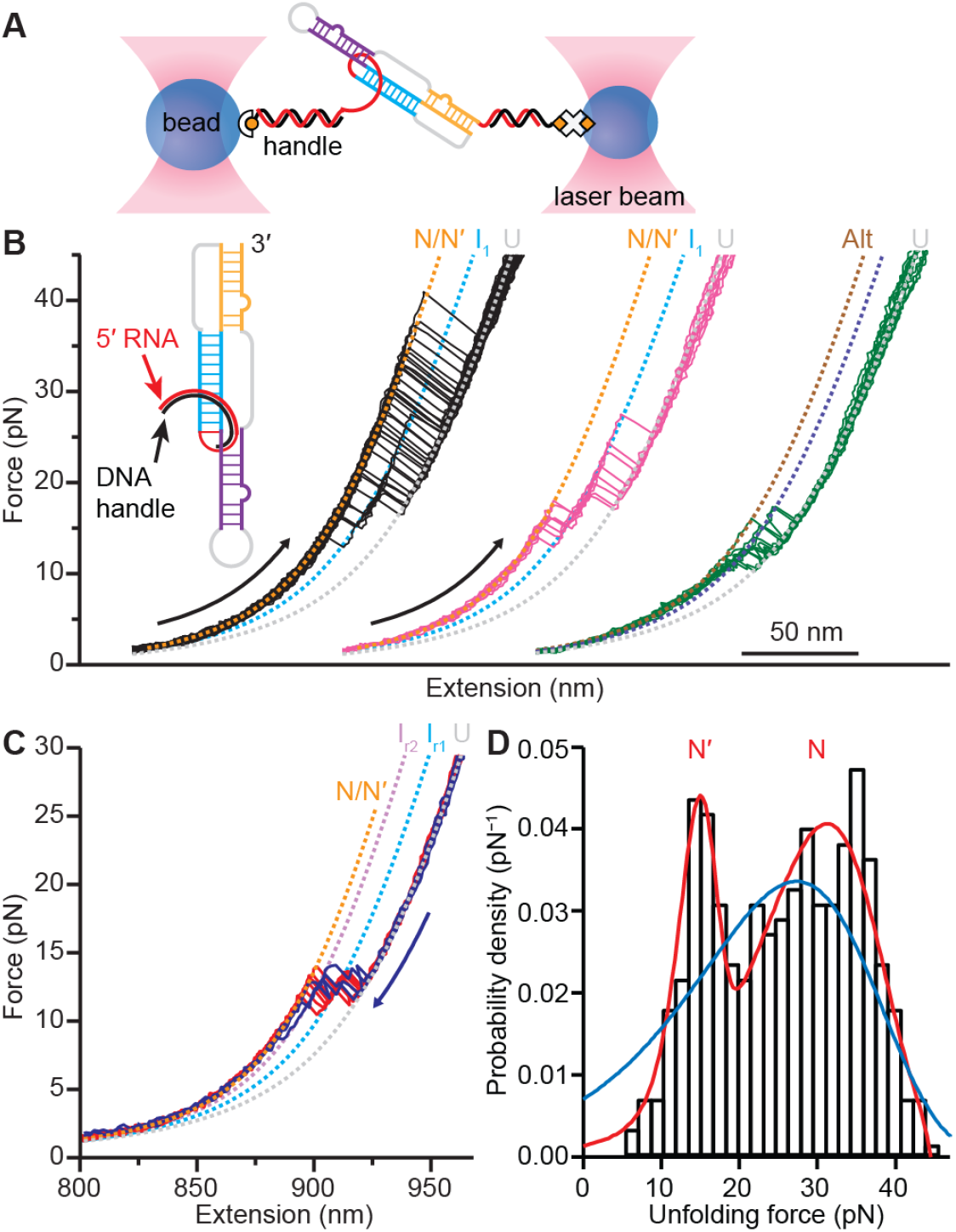
Force spectroscopy of SARS-CoV-2 pseudoknot. (A) Schematic of force spectroscopy assay: a single RNA pseudoknot is tethered by duplex handles attached to beads held in optical traps. (B) Unfolding force-extension curves (FECs) measured with handle edge located 1 nt from 5′ end of stem 1 in pseudoknot. Black: FECs showing full-length unfolding of pseudoknot. Magenta: FECs unfolding pseudoknot via intermediate state I1. Green: FECs showing unfolding of alternate stem-loop structures, including intermediate I_A_. Dashed lines: fits to worm-like chain model. FECs offset for clarity. (C) FECs showing full-length refolding of pseudoknot, with intermediates corresponding to folding of stem 1 (I_r1_, cyan dashed line) and stem 1 + stem 3 (I_r2_, purple dashed line). (D) Unfolding force distribution for pseudoknot shows two peaks, representing two distinct conformers N and N′. Blue: fit to single-population landscape model; red: fit to two-population landscape model.

Examining the unfolding FECs in more detail, we found two qualitatively different behaviors distinguished by different length changes. Measuring the amount of RNA unfolded by fitting the FECs before and after the transition to worm-like chain polymer elasticity models (Eq. 1, Methods) for the handles and unfolded RNA (Fig. 2B, dashed lines), we found that ~80% of FECs (Fig. 2B, black) showed a contour length change of Δ*L*_c_ = 35.1 ± 0.3 nm for complete unfolding (Table S1). This result was close to the value expected for full unfolding of the pseudoknot, 34.7–36.5 nm, based on cryo-EM reconstructions^10,15^ and computational modeling of likely structures^18^. Sometimes these curves contained an intermediate state, I1 (Fig. 2B, magenta), which unfolded with a length corresponding to stem 1 (Δ*L*_c_ = 17.5 ± 0.4 nm), but most of the time they showed a single cooperative unfolding transition. The remaining ~20% of the FECs (Fig. 2B, green) unfolded with a smaller total length change of Δ*L*_c_ = 25 ± 1 nm, indicating an alternate structure that was incompletely folded, at forces of ~10–20 pN; such alternative structures have been observed previously for many frameshift signals^22,33^. This second class of FECs also often contained at least one unfolding intermediate.

There is a characteristic shape expected for the distribution of unfolding forces, *p*(*F*_u_), for unfolding across a single barrier^34^, hence *p*(*F*_u_) can reveal the presence of distinct initial states during the unfolding^35,36^. For the population of FECs with full-length unfolding, two peaks were seen in *p*(*F*_u_): a minor peak near 16 pN and a larger peak near 30 pN (Fig. 2D, black). The double peak indicates the presence of at least two distinct initial conformers, which despite sharing the same total length change nevertheless unfold over different barriers, leading to different shapes for their unfolding force distributions. Such behavior has been seen previously in ligand-bound riboswitches^37^ and proteins^35^. By fitting *p*(*F*_u_) to a kinetic model for barrier crossing (Eq. 2, Methods), the shape of the energy barrier can be characterized through its height (Δ*G*^‡^) and distance from the folded state (Δ*x*^‡^), reporting on the nature of the interactions that hold the structure together^32^. We found that *p*(*F*_u_) did not fit well to the distribution expected for a single initial state (Fig. 2D, blue), but it did fit well to the distribution expected for two initial states (Fig. 2D, red; fit parameters listed in Table S2). The results for Δ*G*^‡^ were similar within error for the two initial states, respectively 40 ± 10 kJ/mol for the higher-force state (denoted N) and 33 ± 8 kJ/mol for the lower-force state (denoted N′), but Δ*x*^‡^ was notably smaller for N: 0.7 ± 0.1 nm, compared to 2.8 ± 0.6 nm for N′, implying a less rigid structure for N′. In both cases, however, the value for Δ*x*^‡^ was consistent with the range characteristic of pseudoknots^22,38^ and other structures containing tertiary contacts^39–41^, but too short for structures consisting only of stem-loops^42^.

Given that the SARS-CoV-2 pseudoknot is predicted to form different fold topologies, such as the 5′-threaded and unthreaded conformers seen in simulations^18^, such different conformers would be expected to give rise to sub-populations with different mechanical properties, because 5′-threaded folds are more mechanically resistant than unthreaded folds^21^. To test if the high-force population involved threading of the 5′ end, we explored if the proportions of the high-force and low-force populations was modulated by changing the proximity of the duplex handle to the 5′ end of stem 1: steric hindrance from a too-close duplex would reduce the likelihood of 5′- end threading. We first re-measured the FECs after moving the duplex 6 nt away from stem 1, instead of 1 nt away as in the measurements in Fig. 2, to make threading easier (Fig. 3A). We found the same contour length changes as before (Table S1), but now the lower-force peak in *p*(*F*_u_) was reduced to a small shoulder (Fig. 3B), with the fraction of FECs showing full-length Δ*L*_c_ attributed to N′ reduced from 20 ± 3% of the FECs to 9 ± 2%, and the fraction attributed to N increased correspondingly from 80 ± 3% to 91 ± 2%. In contrast, when the handle was instead extended past the 5′ end of the pseudoknot so that it paired with the first 2 nts in stem 1 (Fig. 3C, inset), the FECs (Fig. S1) revealed an unfolding force distribution with a significant increase in the occurrence of N′, to 45 ± 4% of the curves with full-length Δ*L*_c_, and a corresponding decrease in N to 55 ± 4% (Fig. 3C). Extending the handle duplex closer to the 5′ end thus produced a clear trend, suppressing N but enhancing N′ (Fig. 3D). However, the occupancy of the alternative, non-pseudoknotted structures was unchanged as the handle was lengthened (Table S1), and the landscape parameters for the two populations also remained the same within error (Table S2). The fact that the only significant effect of changing the handles was to rebalance the ratio of N′ to N supports the conclusion that N is a 5′-threaded conformer, whereas N′ is an unthreaded conformer.

**Fig. 3:**
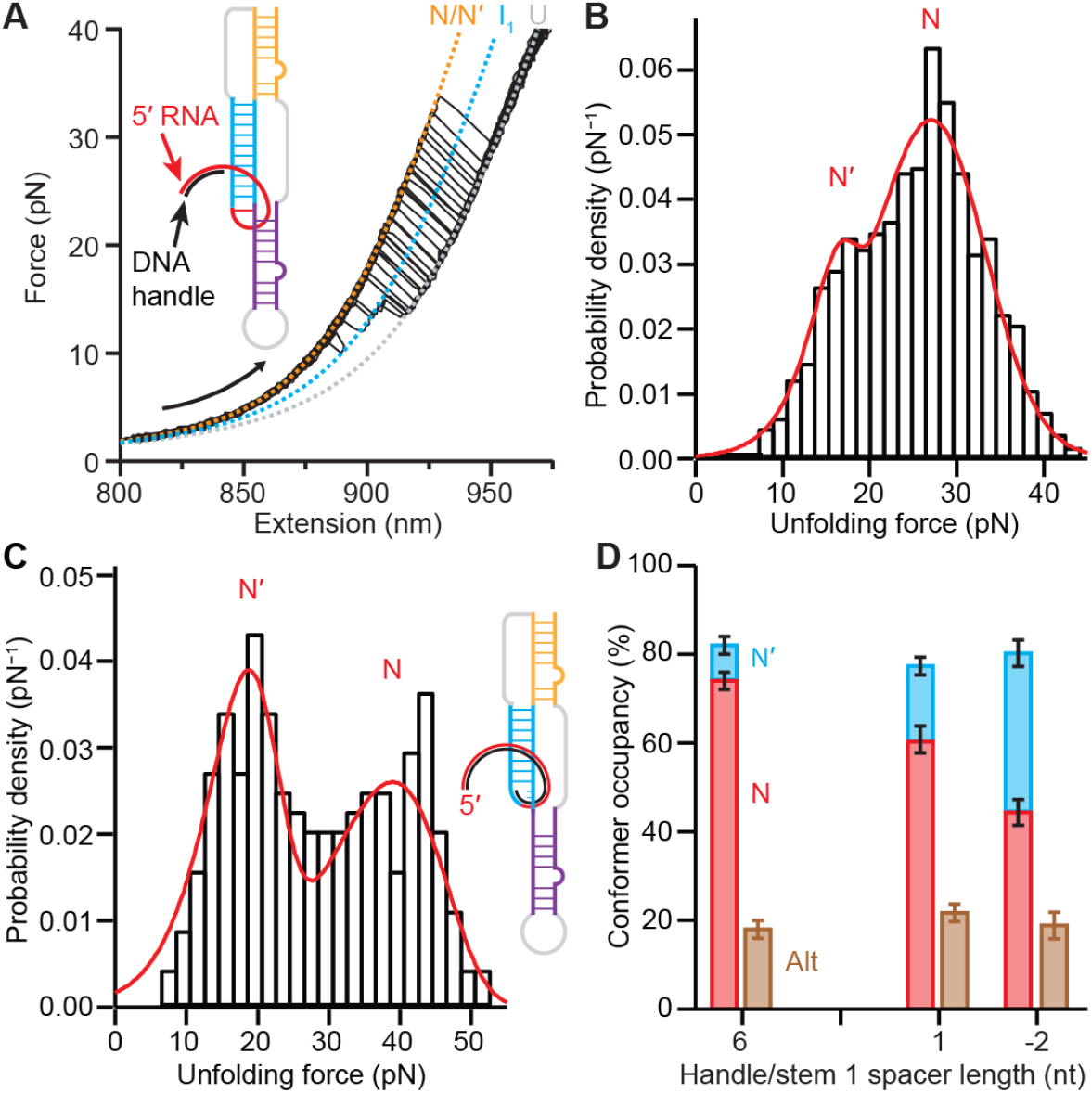
Occlusion of 5′ end alters proportion of pseudoknot conformers. (A) Unfolding FECs measured with edge of duplex handle distanced by 6 nt from 5′ end of pseudoknot show the same length changes as in Fig. 2. (B) Unfolding force distribution with 6-nt spacer between handle and pseudoknot shows less prominent lower-force peak and more prominent higher-force peak. (C) With handle extending 2 nt into stem 1 of pseudoknot (inset), unfolding force distribution shows more prominent lower-force peak and less prominent higher-force peak. (D) Fraction of FECs with higher-force pseudoknot unfolding (red) decreases systematically as duplex handle is extended towards stem 1 of pseudoknot (or invades it), with compensating increase in fraction with lower-force pseudoknot unfolding (cyan), whereas fraction with alternative stem-loop unfolding (brown) remains constant.

Turning to the refolding FECs, we found that the pseudoknotted conformers always refolded through one or more intermediate states. The first refolding transition, which was in the force range ~10–15 pN, had Δ*L*_c_ = 17.7 ± 0.6 nm (Fig. 2C), consistent with the value of 16.7 nm expected for folding of stem 1 in both the threaded and unthreaded models^18^ (Table S1, Fig. S2). After forming stem 1, often the pseudoknot refolded all at once (Fig. 2C, red), but other times it refolded through an additional intermediate (Fig. 2C, blue). The cumulative length changes from the unfolded state for these subsequent transitions were Δ*L*_c_ = 29.2 ± 0.6 nm and 35.3 ± 0.6 nm, consistent with expectations respectively for folding stem 3 as well as stem 1 (29.8 nm), and then stem 2 to form the complete pseudoknot (34.7–36.3 nm for 5′-threaded and unthreaded conformers). Stem 2 thus folded last, after stem 3. Intriguingly, this order is precisely what is needed to form a 5′-threaded fold topology: stem 2 must form last, after the 5′ end is passed over the junction between stems 1 and 3. To confirm the identification of these structures in the intermediates, we repeated the measurements using anti-sense oligos to block the formation either of stem 1 (oligo 1) or stem 2 and part of stem 3 (oligo 2), as shown in Fig. 4A. The initial refolding transitions with oligo 2 present (Fig. 4B) showed effectively the same *p*(*F*_u_) (Fig. 4C, red) as without the oligo (Fig. 4C, black), and close to the same Δ*L*_c_, too, albeit elongated by an extra ~2 base-pairs formed with the part of stem 3 liberated by oligo 2 (Table S1, Fig. S2), confirming that stem 1 was first to refold. FECs with oligo 1 (Fig. 4D), on the other hand, showed refolding at notably lower force than stem 1 (Fig. 4C, blue); unfolding proceeded via an intermediate with a length corresponding to the lower half of stem 3, Δ*L*_c_ = 8 ± 1 nm (Table S1, Fig. S2).

**Fig. 4:**
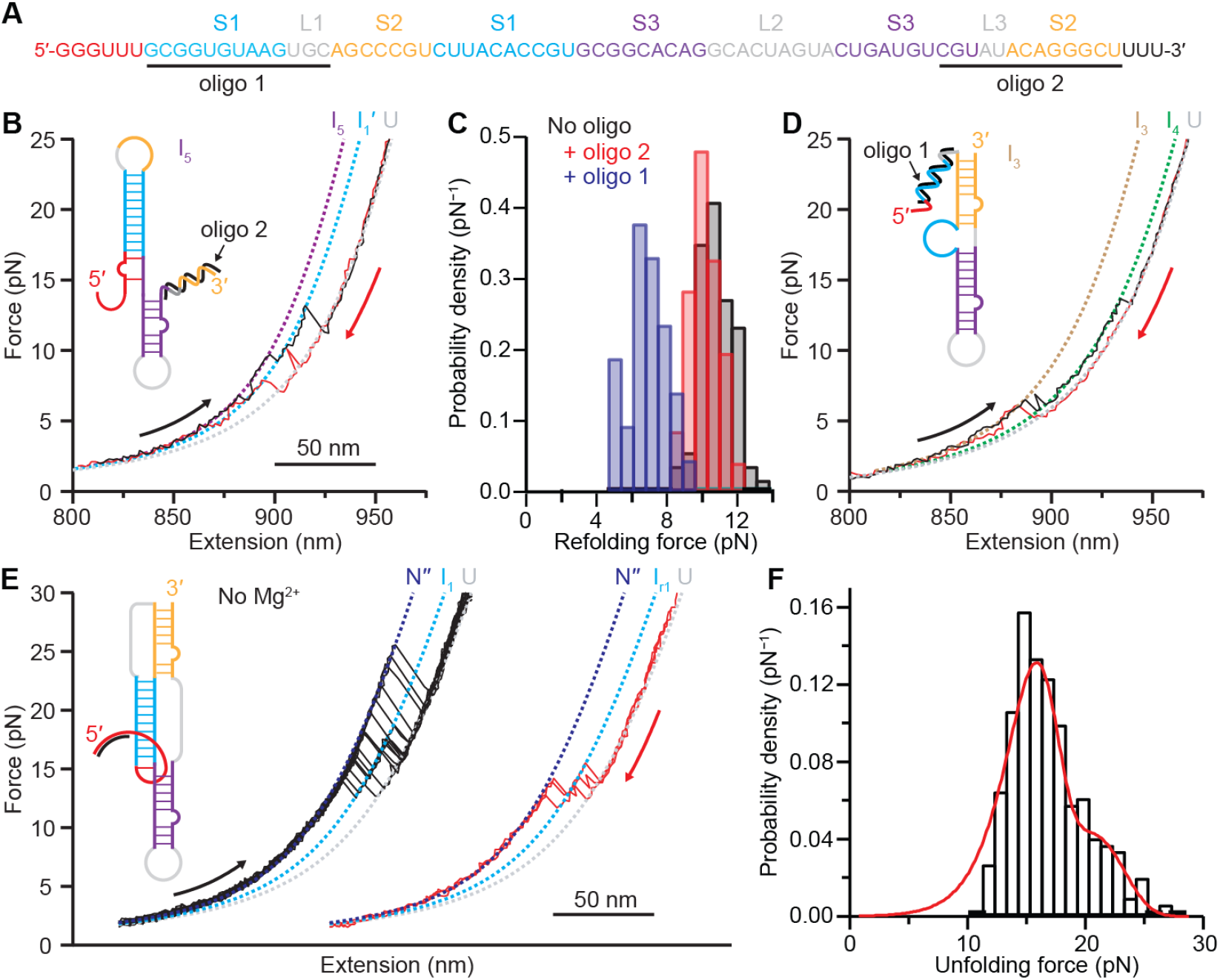
Effects of anti-sense oligomers and Mg^2+^ ions. (A) Anti-sense oligo 1 blocks formation of stem 1, oligo 2 blocks formation of stem 2. (B) FECs with oligo 2 present unfold and refold through an extended stem 1 (I_1_′). (C) Refolding force distribution with oligo 2 present (red) is same as without oligos (black), but higher than with oligo 1 present (blue). (D) FECs with oligo 1 present show folding of stems 2 and 3 but not stem 1, through an intermediate (I_4_) consistent with the lower part of stem 3. (E) FECs without Mg^2+^ show unfolding (black) and refolding (red) of pseudoknotted structures (N′′) similar to FECs with Mg^2+^, through same intermediate I_1_, but with slightly shorter contour length change. Curves offset for clarity. (F) Unfolding force distribution shows two peaks, both at lower force than with Mg^2+^. Red: two-population fit to landscape model.

Finally, we tested the importance of Mg^2+^ for the folding and stability of the pseudoknot by re-measuring FECs in the absence of Mg^2+^ (Fig. 4E). We found that 97% of curves showed the length change for pseudoknot unfolding, although Δ*L*_c_ was ~ 1 nm shorter than previously (Table S1). Two populations were still present in *p*(*F*_u_) (Fig. 4F), but the high-force population was greatly reduced (down to only 40 ± 8% of the FECs showing full-length unfolding), and it peaked at lower forces. Mg^2+^ was therefore not required for folding of the pseudoknot, but it played a key role in promoting the formation of the higher-force population attributed to the ring-knot fold topology. Fitting *p*(*F*_u_) to characterize changes in the landscape (Fig. 4F, red), we found that Δ*G*^‡^ was little changed, but Δ*x*^‡^ was significantly higher, rising to 4.0 ± 0.6 nm for N′ and 2.8 ± 0.7 nm for N (Table S2), indicating that the pseudoknots were much less rigid without Mg^2+^.

## Discussion

These results confirm the suggestion from simulations and cryo-EM imaging that the SARS-CoV-2 frameshift signal can form a variety of different structures, and they reveal how these structures respond to mechanical tension similar to what the ribosome would apply during −1 PRF. The most prominent conformation, N, unfolded through the full length of the pseudoknot at moderately high force and was suppressed by occlusion of the 5′ end by the duplex handle, precisely as would be expected for a 5′-end threaded structure such as those seen in cryo-EM images on and off the ribosome^10,15^ or predicted from simulations^18^. In contrast, the conformation unfolding at lower force (N′) occurred more frequently when the 5′ end was occluded, in proportion to the suppression of N, as would be expected for a conformation in which the 5′ end remains unthreaded. Although predicted computationally^18^, unthreaded conformations have not yet been identified in cryo-EM images; they are nevertheless expected to be present at some level because they will form naturally in a kinetic partitioning process whenever stem 2 folds while the 5′ end is not lying across the stem 1/stem 3 junction (as required for 5′-end threading), similar to what was seen in the folding of the Zika exonuclease-resistant RNA^21^.

The picture that emerges of the pseudoknot folding and unfolding is illustrated in Fig. 5. Stem 1 always folds first, followed either by sequential folding of stem 3 then stem 2, or else cooperative refolding of both stems simultaneously. The orientation of the 5′ end at the moment of stem 2 formation partitions the molecule into two distinct fold topologies that cannot interconvert: 5′-threaded or unthreaded. These two fold topologies then give rise to distinct unfolding behaviors: higher forces for the threaded fold, lower forces for the unthreaded fold. Notably absent from this picture, however, is a third fold that was predicted computationally with the 3′ threaded through the stem 2/stem 3 junction^18^, given that this fold requires stem 1 to form last instead of first.

**Fig. 5:**
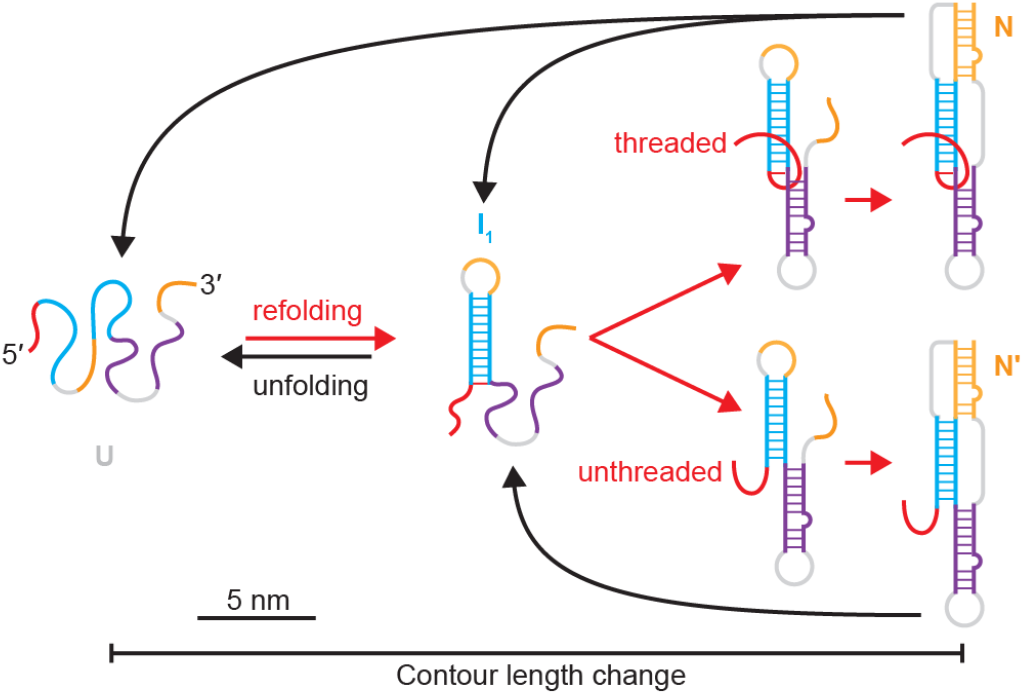
Pseudoknot folding and unfolding pathways. Schematic of pathways for folding (red) and unfolding (black) the pseudoknotted conformers. Stem 1 forms first, then stem 3, leading to either 5′-threaded (top) or unthreaded (bottom) conformers depending on orientation of 5′ end just before stem 2 forms.

The energy landscape parameters for unfolding states N and N′ further support this picture, in particular Δ*x*^‡^, which reports on the mechanical rigidity of the RNA structure. Smaller Δ*x*^‡^ implies a rigid structure that is less sensitive to tension and hence more likely rupture in a brittle manner at high force, whereas larger Δ*x*^‡^ implies a compliant structure that is more sensitive to tension and ruptures in a lower, narrower range of forces. State N was much more mechanically rigid than N′, with Δ*x*^‡^ roughly 3 times smaller, which makes sense in terms of the mechanical effects of threading: threading of the 5′ end should rigidify the fold via interactions between the 5′ end and the helical junction in the pseudoknot^18^ that constrain the motion of the terminus in response to tension applied to the 5′ end, compared to the unthreaded fold. Indeed, the Δ*x*^‡^ value of only 0.7 nm for state N is among the smallest reported for any pseudoknot, signifying the particular rigidity of this unusual fold topology; in contrast, the Δ*x*^‡^ value for N′ is comparable to the range of values more typically reported for other pseudoknots, ~1.5–2 nm^22,38^. The fact that Δ*x*^‡^ increased significantly for both N and N′ in the absence of Mg^2+^ shows that although Mg^2+^ is not required for the pseudoknot folding, consistent with results from NMR^16^, it is essential for stabilizing tertiary contacts that rigidify both the threaded and unthreaded folds. Comparing the NMR results to computational modeling^18^ suggests that Mg^2+^-mediated tertiary interactions may be especially important in stem 2, explaining why both N and N′ are more compliant in the absence of Mg^2+^.

Comparing the SARS-CoV-2 pseudoknot to the Zika virus exonuclease-resistant RNA (xrRNA), the only other RNA forming a similar ring-knot whose folding has been studied, reveals some interesting differences. The ring-knot in the Zika virus xrRNA unfolds at a much higher force, above 60 pN, acting as a mechanical road-block to digestion of the viral RNA by host exo-ribonucleases^21^. Such extreme mechanical resistance would be functionally counterproductive for the SARS-CoV-2 pseudoknot, however, which acts to induce a frameshift but does not prevent ribosomal translocation. Concurrently, these two RNAs differ in the importance of Mg^2+^ for threading of the 5′ end before closure of the pseudoknot: threading is abolished entirely for the Zika virus xrRNA in the absence of Mg^2+^, but is only partially inhibited (not abolished) for the SARS-CoV-2 pseudoknot, indicating that at least some of the interactions that stablize threading must be independent of Mg^2+^. We speculate that the reduced role of Mg^2+^ in 5′-threading in the SARS-CoV-2 pseudoknot may reduce the mechanical stability of the ring-knot sufficiently to allow the ribosome to unfold it during −1 PRF.

Finally, we note that the significant heterogeneity seen for the SARS-CoV-2 frameshift signal is entirely consistent with the direct correlation between conformational heterogeneity and −1 PRF efficiency found in recent work^27^, given the relatively high level of −1 PRF in SARS-CoV-2 observed in functional assays^1,11^. The existence of distinct fold topologies also has important implications for structure-based drug-discovery efforts targeting the SARS-CoV-2 frameshift signal, because the structure of the junction between the helices in the pseudoknot, which is the locus of the most likely binding pockets for small molecules^14,43^, is strongly affected by whether the 5′ end is threaded or not^18^. Combining these two observations suggests a strategy for developing small-molecule modulators of −1 PRF in SARS-CoV-2: since −1 PRF efficiency varies directly with the conformational Shannon entropy^27^, ligands that stabilize the 5′-threaded conformer (thereby decreasing the heterogeneity) should be sought for inhibiting −1 PRF, whereas ligands that stabilize the unthreaded conformer (thereby increasing the heterogeneity) should be sought for enhancing −1 PRF.

## Methods

### Sample preparation

Samples consisting of a single RNA strand linked at each end to double-stranded handles were prepared in two ways. (1) An RNA strand containing the SARS-CoV-2 pseudoknot and spacer sequences (Fig 1A) flanked by long ‘handle’ sequences was annealed to single-stranded (ss) DNA complementary to the handles sequences, as described previously^22^. The DNA fragment corresponding to the sequence in Fig. 1A was cloned into the pMLuc-1 plasmid between the BamHI and SpeI sites. A 2,749-bp DNA transcription template was amplified by PCR from this plasmid, containing a T7 promoter in the upstream primer, followed by a 1,882-bp handle sequence, the pseudoknot in the middle, and then a 798-bp handle sequence downstream; RNA was transcribed from this template in vitro. Two ssDNA handles complementary to the upstream and downstream handle sequences in the RNA were created by asymmetric PCR. The 3′ end of the 1,882-nt DNA handle was labeled with dig-ddUTP using terminal transferase (Roche), and the 798-nt DNA handle of the transcript was functionalized with biotin on the 5′ end of the PCR primer. The RNA transcript was annealed to the DNA handles. (2) A shorter RNA strand was annealed to a 300-nt ssDNA handle on one end, and to the overhang on a 2,094-bp double-stranded (ds) DNA handle on the other end. The dsDNA handle, labeled with digoxigenin via the upstream primer, was prepared by digesting a 2,075-bp PCR product amplified from the plasmid pUC19 with PspGI, and then ligating a 56-nt DNA oligo to the digest sticky end to create a 36-nt 3′ overhang. RNA was transcribed in vitro from a 406-bp DNA template containing a T7 promoter, the 36-nt sequence complementary to the ssDNA overhang, the pseudoknot and spacer sequences (Fig. 1A), and a 300-nt handle sequence; this transcription template was made by cloning the required DNA sequences into a modified pMLuc-1 vector between the XhoI and SpeI sites. A 300-nt ssDNA handle complementary to the handle region of the RNA transcript was made by asymmetric PCR and annealed to the RNA, as above. The resulting DNA-RNA complex was then annealed to the overhang of the dsDNA handle.

Constructs were readied for measurement as described previously^22^. Briefly, the RNA/handle constructs were diluted to ~160 pM and mixed with equal volumes of 600-nm and 820-nm diameter polystyrene beads (coated respectively with avidin DN and anti-digoxigenin) at concentrations of ~250 pM, and incubated for ~1 hour at room temperature to create ‘dumbbells’. The incubation was then diluted ~100-fold into RNAse-free measuring buffer: 50 mM MOPS pH 7.5, 130 mM KCl, 4mM MgCl2, and 200 U/mL RNAse inhibitor (SUPERase•In, Ambion). An oxygen scavenging system consisting of 40 U/mL glucose oxidase, 185 U/mL catalase, and 250 mM D-glucose was also included in the buffer. The diluted dumbbells were placed in a flow chamber prepared on a microscope slide and inserted into the optical trap.

### FEC measurements and data analysis

Measurements were made using custom-built optical traps described previously^37^. Traps were calibrated for position detection for each dumbbell prior to measurement, as described previously^44^. FECs were measured by moving the traps apart at a constant speed of ~160 nm/s to increase the force up to ~50 pN and unfold the RNA, bringing them back together at the same speed to ramp the force back down to ~0 pN, waiting 5–10 s to allow refolding, and then repeating the cycle. Trap stiffnesses were 0.45–0.62 pN/nm. Data were sampled at 20 kHz and filtered online at the Nyquist frequency. Measurements with anti-sense oligos added 10 μM oligo 1 or oligo 2 to the measuring buffer. For measurements in the absence of Mg^2+^, we removed the MgCl2 from the measuring buffer and added 1 mM EDTA.

Each of the branches of the FECs separated by ‘rips’ representing unfolding/refolding transitions was fit to an extensible worm-like chain (WLC) model relating the applied force, *F*, and molecular extension, *x*:

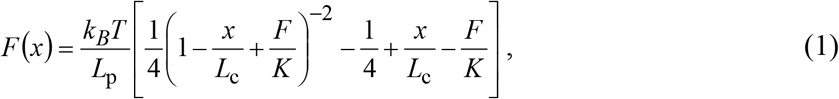

where *L*_p_ is the persistence length, *L*_c_ the contour length, and *K* the enthalpic elasticity^45^. Two WLCs in series were used for the fitting, one to describe the duplex handles and the other for the unfolded RNA^37^. The WLC parameters for the handles, found from fitting the folded state of the FECs, were typically *L*_p_ ~ 40 nm, *L*_c_ ~ 850 nm for the shorter construct and ~950 nm for the longer one, and *K* ~ 1000 pN. The parameters for the unfolded RNA were fixed at *L*_p_ = 1 nm, *L*_c_ = 0.59 nm/nt, and *K* = 1500 pN, so that the only free parameter was the number of nucleotides unfolded.

Force distributions were fit to the theory of Dudko et al^34^:

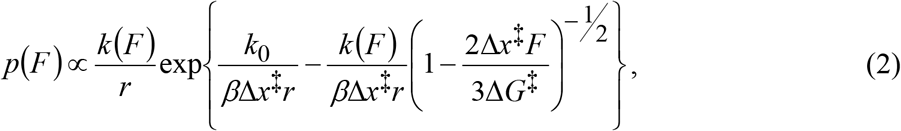

where 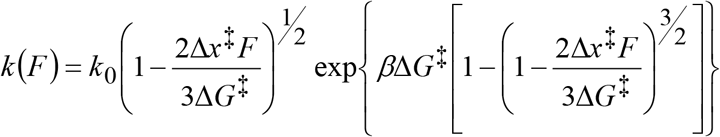, *k*_0_ is the unfolding rate at zero force, Δ*x*^‡^ is the distance from the folded state to the barrier, Δ*G*^‡^ is the barrier height, and 1/*β* = *k*B*T* is the thermal energy. Distributions of the unfolding forces showing two peaks were fit by a sum of two independent distributions represented by Eq. 2. Errors for the fitting parameters were found from bootstrapping analysis. We verified that two-population fits of *p*(*F*_u_) were justified using the Akaike information criterion to test for overfitting and the Wald-Wolfowitz runs test to assess underfitting. These tests rejected the single-population fits in favor of the double-population fits in each case.

## Acknowledgements

This work was supported by the Canadian Institutes of Health Research, Alberta Innovates, and National Research Council Canada.

## Supporting information

**Table S1:**
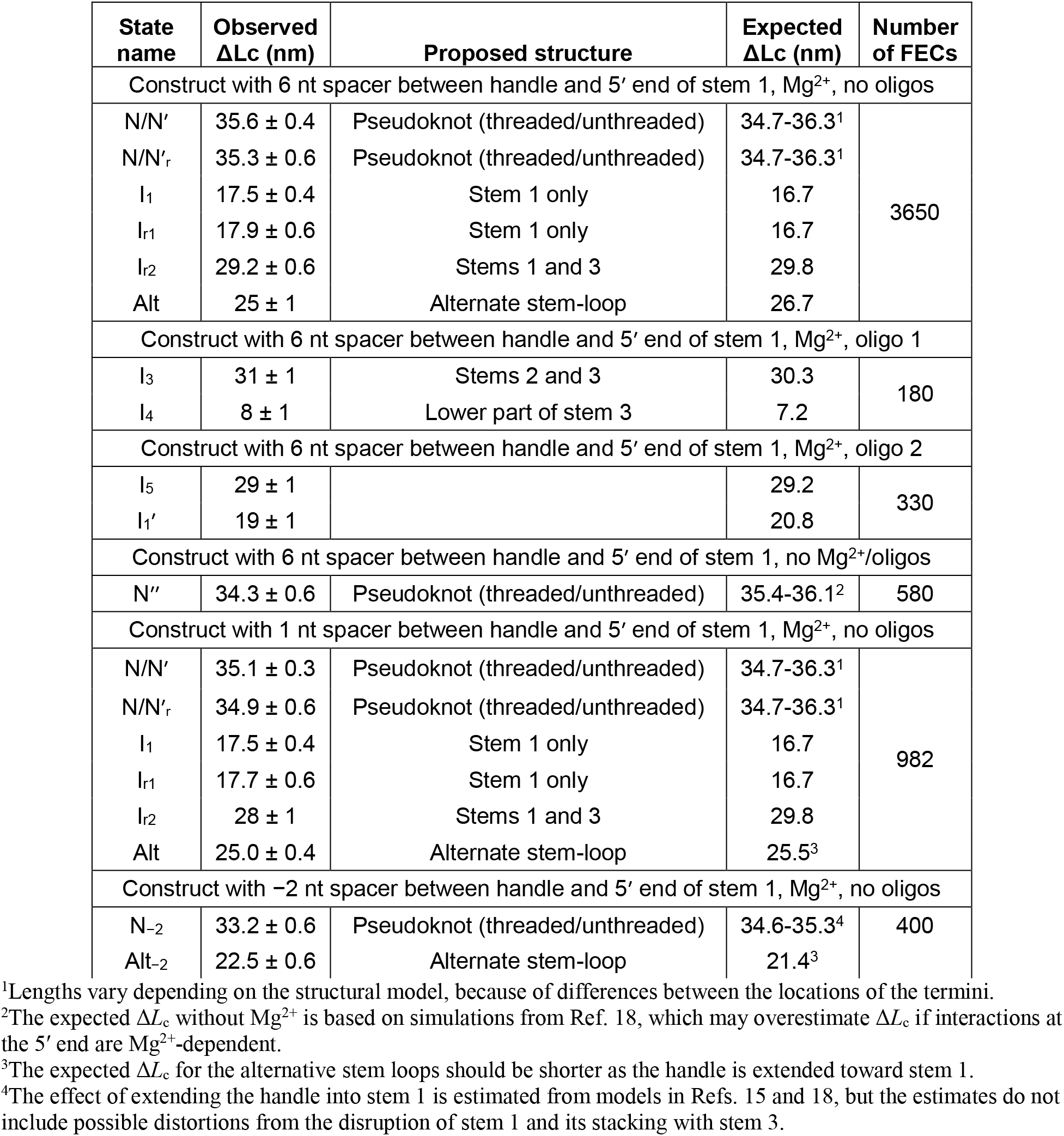
Length changes for unfolding and refolding transitions. State names are shown in FECs in Figs. 2–4 and Fig. S2. Subscript “r” indicates results from refolding. Expected values are based on the structural models shown in Fig. S2. Errors on measured values represent standard error on the mean.

**Table S2:** Energy landscape fit parameters. Errors represent standard error on the mean from bootstrapping analysis.

| Spacer length between handle and stem 1 (nt): | Lower-force population |  |  | Higher-force population |  |  |
| --- | --- | --- | --- | --- | --- | --- |
| | $\ln(k_0)$ [s<sup>-1</sup>] | $\Delta x^\ddagger$(nm) | $\Delta G^\ddagger$(kJ/mol) | $\ln(k_0)$ [s<sup>-1</sup>] | $\Delta x^\ddagger$(nm) | $\Delta G^\ddagger$(kJ/mol) |
| 6 | -5 ± 1 | 2.1 ± 0.3 | 33 ± 5 | -4 ± 1 | 0.7 ± 0.1 | 31 ± 4 |
| 1 | -6 ± 1 | 2.8 ± 0.6 | 33 ± 8 | -3.5 ± 0.3 | 0.7 ± 0.1 | 40 ± 10 |
| -2 (invades stem 1) | -2.8 ± 0.4 | 1.5 ± 0.2 | 27 ± 6 | -5.4 ± 0.6 | 0.7 ± 0.1 | 29 ± 6 |
| 6 (without Mg<sup>2+</sup>) | -8 ± 1 | 4.0 ± 0.6 | 40 ± 6 | -7 ± 1 | 2.8 ± 0.7 | 35 ± 6 |

**Fig. S1:**
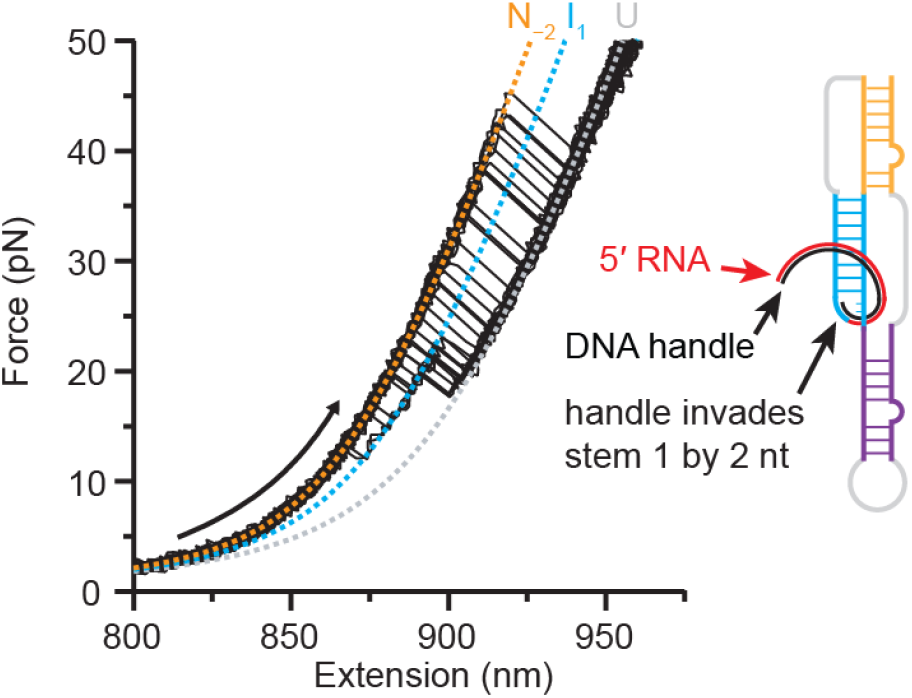
FECs measured with handle extending 2 nt into stem 1. Unfolding FECs show the same qualitative behavior seen in Fig. 2 and Fig. 3A, except that the total length change is ~2 nm shorter and there are significantly more unfolding events at lower force.

**Fig. S2:**
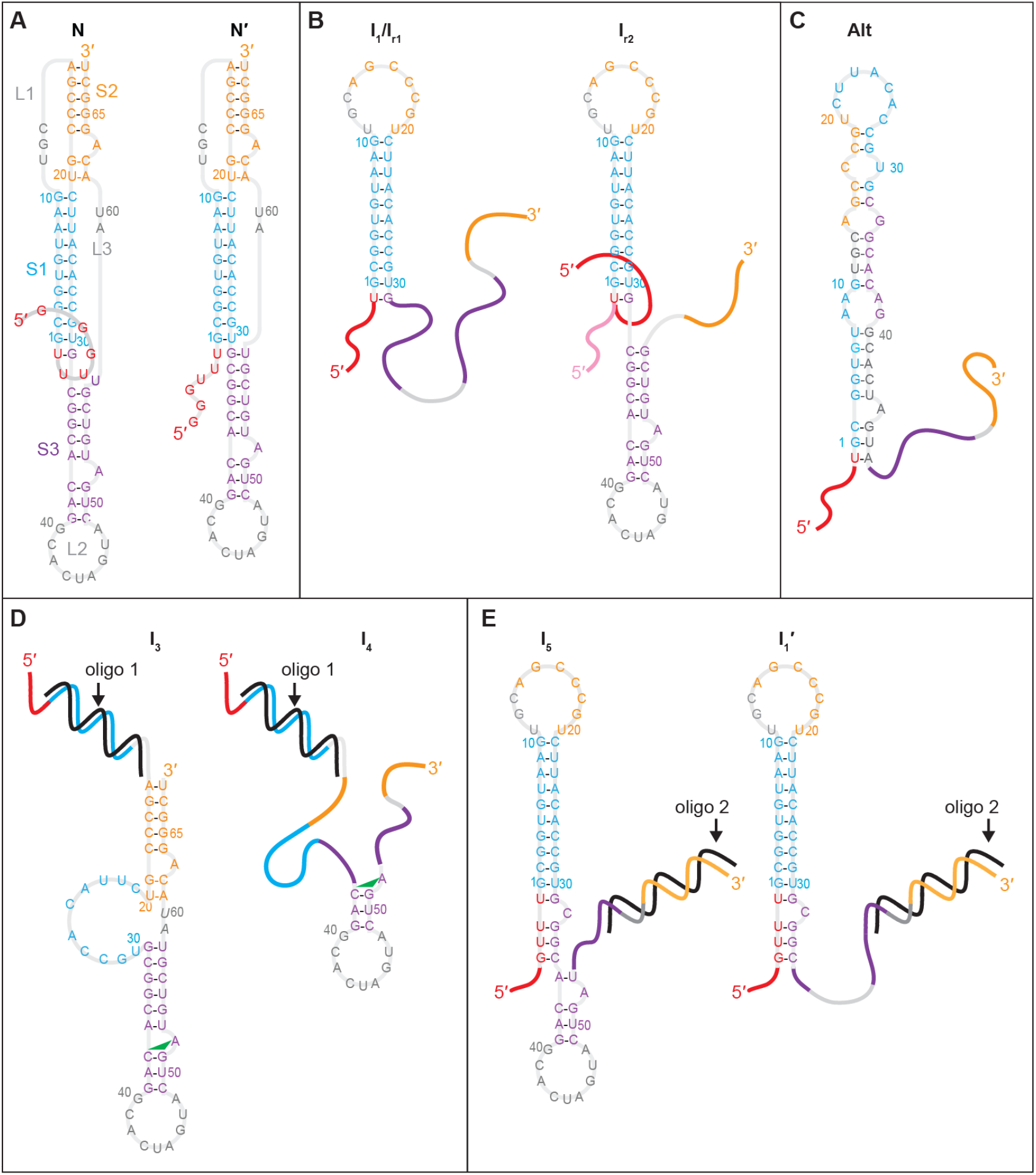
Models of intermediate structures observed in FECs. (A) 5′-threaded (left) and unthreaded (right) conformers of pseudoknot, as per models in Ref. 18. (B) Unfolding and refolding intermediates with stem 1 only (left) and both stems 1 and 3 (right). For I_r2_, the geometry of the 5′ end just before closure of stem 2 determines the fold topology: threaded (red) or unthreaded (pink). (C) Model of alternative stem-loop structure, which prevents pseudoknot formation. (D) Structures formed with oligo 1 bound to stem 1. Intermediate I_4_ is proposed to be stabilized by the C37:G51:A52 triple (green triangle) found in Ref. 18. (E) Structures formed with oligo 2 bound to stem 2. Stem 1 is extended to pair with some of the unpaired part of stem 3.

